# Long-term population dynamics of an endangered butterfly are influenced by hurricane-mediated disturbance

**DOI:** 10.1101/2024.07.12.603220

**Authors:** Sarah R. Steele Cabrera, Michael Belitz, Thomas C. Emmel, Emily S. Khazan, Matthew J. Standridge, Kristin Rossetti, Jaret C. Daniels

## Abstract

Effective species conservation requires understanding an organism’s population dynamics and natural history, but long-term data are challenging to collect and maintain. As a result, conservation management decisions are frequently made using short-term data, which are insufficient to accurately assess population trends in most species. For less-studied taxa, including most invertebrates, inadequate understanding of life and natural history also impedes conservation efforts. Long-term studies are highly valuable for improving conservation decisions for target species as they serve as a model for other understudied species. We use mark-recapture data collected over 35 years to examine weather drivers of population patterns for an endangered butterfly, Schaus’ swallowtail (*Heraclides ponceana*), and to enhance our understanding of its natural history. We show that the population size of Schaus’ swallowtail butterfly was highly variable, ranging from under 100 to over 10,000 individuals. Population size is influenced by weather events and population size in the previous year. Population size was lower immediately following high wind events but was positively influenced by high wind events four years prior, with notable population increases following tropical cyclone events. Precipitation during the dry season preceding the adult flight period was also associated with higher population sizes. This study reveals the potentially beneficial role of hurricane-mediated disturbance on Schaus’ swallowtail populations potentially due to increased treefall gaps and the resulting shifts in plant communities. This remarkable data set represents one of the longest-term studies on a tropical insect.

## 1. Introduction

Understanding the long-term population dynamics and natural history of organisms is fundamental in ecology (Magurran et al. 2010; Siddig et al. 2016). In the face of global climate change and the current biodiversity crisis (Manes et al. 2021; McLaughlin et al. 2022), this information is fundamental to biodiversity conservation, especially for species which are already vulnerable to extinction (Nielsen et al. 2009). One conservation challenge, particularly for invertebrates, is a lack of understanding of the natural history of organisms that are under threat of extinction (Cardoso et al. 2011; Hochkirch et al. 2021). Conservation of organisms with complex life histories, such as insects, requires comprehensive understanding of their natural history since different life stages often have disparate ecological niches (Wilbur 1980; Nakazawa 2015).

Partial information about species life and natural history, for example knowledge of only one life stage (e.g. adult butterflies) leads to misinformed conservation decisions. In the case of the large blue butterfly (*Maculinea arion*) in Europe, for example, conservationists inadvertently excluded herbivores which are important to maintaining suitable habitat for the butterfly’s obligate ant host when they erected fences to keep out human collectors of the rare butterfly (Thomas 1980). The importance of herbivores was only revealed decades later through intensive ecological studies; this discovery was central to the success of reintroduced populations of this butterfly in Great Britain (Thomas et al. 2009). Similarly, conservation of St. Francis satyr butterfly (*Neonympha mitchellii francisci*) in North Carolina, USA, was impeded by a lack of understanding of the important role that habitat disturbance by fire played in the persistence of populations, which was revealed through long-term studies (Haddad 2018). Both cases illustrate the importance of a deep understanding of natural history and the role of ecological disturbance in species conservation.

The ecological role of disturbance in the population dynamics of many organisms is notable, especially those which are adapted to and benefit from ecological disturbances such as fire, wind, or flooding (Hobbs and Huenneke 1992). Insects are often especially sensitive to impacts from disturbance (Brown 1997; Schowalter 2012), including those deriving from climate change such as rising temperatures, changes in precipitation patterns, and the increasing frequency and severity of extreme weather events (Nielsen and Papaj 2015; Forrest 2016; Renner and Zohner 2018). Due to their life history and distinct dependence on plants throughout their life cycle, butterflies represent a group of insects particularly sensitive to and reliant upon ecological disturbances such as fire (Mason et al. 2021) and grazing (Vogel et al. 2007). Insect disturbance ecology is best understood for temperate taxa, but largely unknown for most tropical taxa despite the high diversity of tropical insects (Lamarre et al. 2020; Slade and Ong 2023).

Tropical cyclones are an ecologically important form of disturbance that are projected to increase in severity in the Atlantic Basin due to anthropogenic climate change (Bender et al. 2010; Knutson et al. 2010; Walsh et al. 2016). The impact of tropical cyclones on insect communities is highly variable. This is particularly evident in coastal ecosystems which are also vulnerable to disturbances including habitat loss and sea level rise (Henry et al. 2020; Rippel et al. 2021). The influence of tropical cyclones on insect communities is well-studied at Luquillo Experimental Forest in the Luquillo Mountains of eastern Puerto Rico, USA, where experiments on the role of disturbance from hurricanes as well as experimental forest gap creation have been ongoing since the 1980s (Shiels et al. 2015). Studies in this system demonstrate that long-term population dynamics of insects are determined largely by hurricane-mediated disturbance and subsequent forest succession (Shiels et al. 2015; Schowalter et al.

2021; Pandey and Schowalter 2022). Other studies of insect communities following tropical cyclones corroborate findings from Luquillo; increases in insect abundance and diversity following tropical cyclones likely occur due to increased heterogeneity in forest structure and resulting increases in resources (e.g., new plant growth, increased floral abundance) (Mullany et al. 2018; Novais et al. 2018; Badon et al. 2022).

Southern Florida, USA, contains a mixture of Caribbean and continental North American flora and fauna and is one of few subtropical ecological communities in the continental United States (Myers and Ewel 1990), with many plant and insect species colonizing South Florida from Cuba and the Bahamas (Peck 1989; Gillespie 2006). The Florida Keys, an archipelago located off the coast of southern Florida (Figure 1A), comprises part of the Caribbean biodiversity hotspot, which has a high rate of endemism in a relatively small geographic area with high rates of habitat destruction and land use change (Myers et al. 2000). South Florida experienced dramatic land use change in the 20th century, with deforestation of upland habitats concentrated along the coast due to massive human population increases (Walker et al. 1997). The area also experiences regular impacts from tropical cyclones with a major hurricane occurring approximately every 30 years in this region, with less severe storm events occurring in most years (Doyle and Girod 1997).

**Figure 1.**
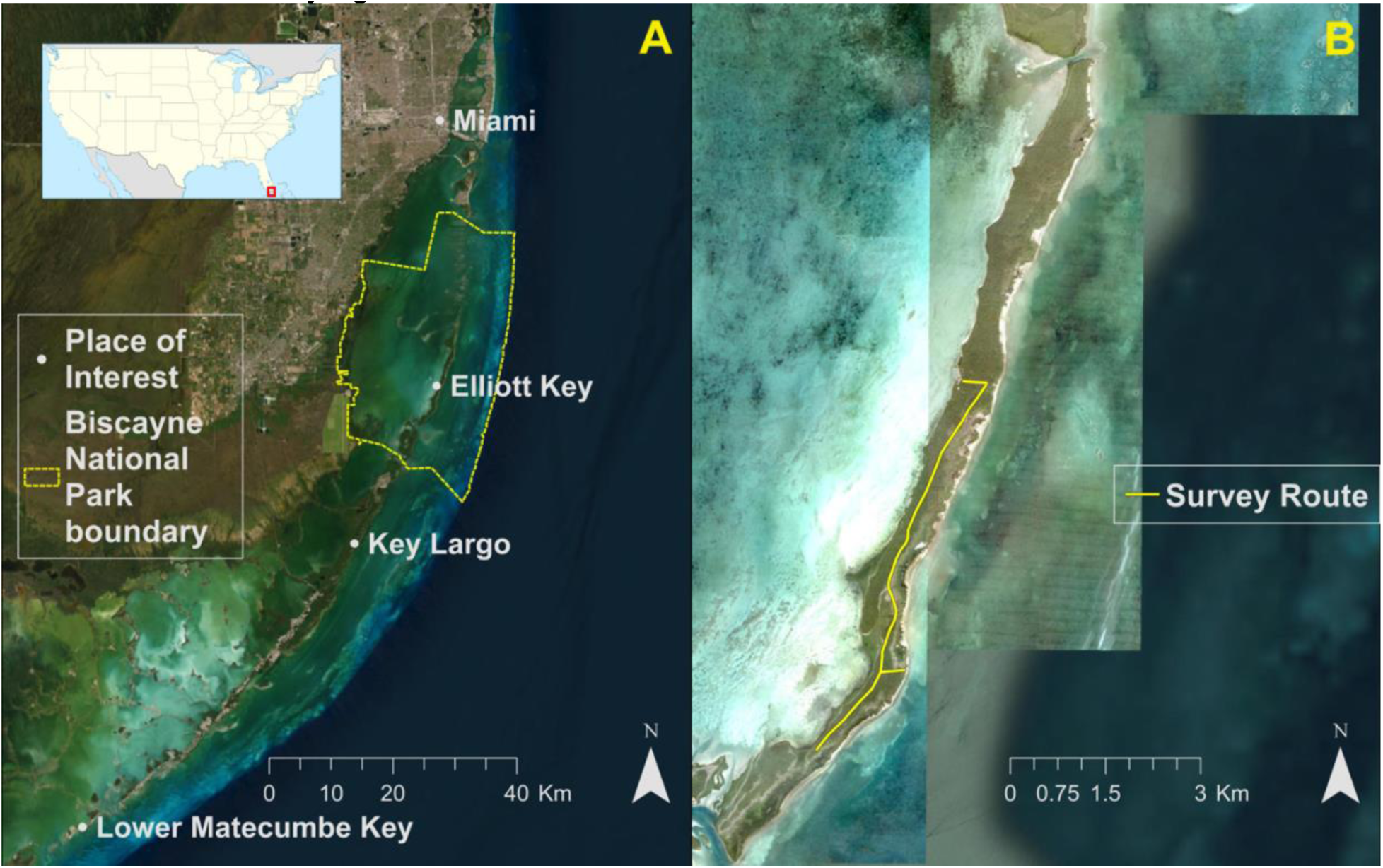
(A) Map showing the historic range of Schaus’ swallowtail butterfly. Schaus’ swallowtail historically occupied hardwood hammocks from Miami to Lower Matecumbe Key but is only extant in northern Key Largo and several islands within Biscayne National Park. (B) Map showing the study site, Elliott Key and the linear survey route which covers much of the butterfly’s habitat on the island. Imagery courtesy Esri and Wikimedia Commons.

Given the biogeographical context and high human development pressure in South Florida, it is perhaps unsurprising that many of the endemic species in the region’s forests are imperiled. These include charismatic mammals such as the Florida panther (*Puma concolor coryi*) and the Key deer (*Odocoileus virginianus clavium*), but also dozens of other species including plants, reptiles, small mammals, and invertebrates (Florida Natural Areas Inventory 2010). Most endemic species in this region are dependent upon scarce freshwater resources and upland forests, including several insects listed as federally endangered: Bartram’s scrub-hairstreak butterfly (*Strymon acis bartrami*) (Emmel and Minno 1993), Florida leafwing butterfly (*Anaea troglodyta floridalis*) (Minno and Emmel 1993), Miami tiger beetle (*Cicindelidia floridana*) (Brzoska et al. 2011), and Schaus’ swallowtail butterfly (*Heraclides ponceana*) (Minno and Emmel 1993).

In this study, we examine the population dynamics of a narrow-range endemic butterfly in the Florida Keys, USA, Schaus’ swallowtail butterfly (*Heraclides ponceana* Schaus 1911). Using intensive monitoring data collected over a 35-year-long period, we explore possible weather drivers of these patterns and aim to better understand the natural history of this butterfly. This study is likely the longest-term intensive monitoring effort for an imperiled tropical butterfly and is an apt study system for research on tropical cyclone disturbance on insect populations. These data are also central to informing ongoing conservation and recovery efforts for this endangered butterfly (U.S. Fish and Wildlife Service (USFWS) (USFWS 2019).

## 2. Material and Methods

### 2.1 Study Species

Schaus’ swallowtail butterfly is a federally endangered papilionid endemic to extreme southeastern Florida (USFWS 1984, 2021). This species was once found from Miami, Florida to Lower Matecumbe Key, Florida (Figure 1A). Since the extirpation of many populations in the early-mid 20th century due to habitat destruction and impacts from insecticides used for mosquito control (USFWS 1984; Eliazar and Emmel 1991), current extant populations are only found within the conservation lands of Biscayne National Park and Key Largo (USFWS 2021). The butterfly is found in hardwood hammock forests, a type of seasonally dry subtropical forest which contain this species’ larval host plants, *Amyris elemifera* L. and *Zanthoxylum fagara* (L.) Sarg., both of which are understory trees in the family Rutaceae. Hardwood hammock forests occur at the highest elevations in the Florida Keys, areas that are most desirable for human development. This has resulted in the loss of most upland habitat across the butterfly’s range (Karim and Main 2009).

Schaus’ swallowtail has one, or possibly two broods per year, with the greatest number of adults present in May and June, corresponding with the beginning of the wet season in South Florida (Loftus and Kushlan 1984; Glassberg et al. 2000). While adults have been observed in August and September, it is unknown whether this represents a true second brood or sporadic asynchronous termination of diapause (Covell 1977; Loftus and Kushlan 1984). The species has been documented to remain in pupal diapause for up to two years in field conditions (Loftus and Kushlan 1984) and even longer in captivity under laboratory conditions (Daniels, unpublished data). Pupae also appear to be robust to sea water flooding and other seemingly detrimental impacts of disturbance (National Park Service (NPS) 2019).

### 2.2 Study Site

Elliott Key (25.433, -80.202; Figure 1B) is the largest island within Biscayne National Park; it is approximately 12 km long and 0.75 km wide and is composed of fossilized coral. Average elevation above sea level on Elliott Key is about 3 meters, and maximum elevation is approximately 5 meters (Klein 1970). Several indigenous groups were known to inhabit this area by 8000 B.C.; the island was later settled in the 19th century for pineapple cultivation (Brown Leynes and Cullison 1998). A six-lane highway was planned for Elliott Key in the 1960s for which a seven-mile-long swath of forest was cleared. Further construction was halted in 1968 with the establishment of Biscayne National Monument, now Biscayne National Park (NPS 2017). This remaining linear clearing is known as “Spite Highway” which is maintained by the NPS and serves as the transect for surveying Schaus’ swallowtail butterflies.

### 2.3 Mark-Release-Recapture study

Surveys were conducted by trained observers along a 6 km section of trail (Figure 1B) between 9:00 and 17:00, when butterflies were observed to be most active. Survey parties consisted of between one and four people traveling either on foot or in an all-terrain vehicle along the transect searching for, capturing, and marking butterflies. For each survey, the most experienced surveyor was designated the primary observer. In each year, surveys were repeatedly conducted between April and June, the main flight period for the butterfly. In some years, additional surveys were conducted in August and September following new observations of butterflies during those months, though the number of observations was too small to include in this analysis. Surveys were not conducted during inclement weather. Beginning in 1986, researchers have produced 23 years of mark-release-recapture data with a median of 10 survey days per year (range from 3 - 29 survey days per year). Some years have missing or low-quality data due to factors including equipment failure, personnel availability, and the COVID-19 pandemic.

Observers attempted to capture every Schaus’ swallowtail butterfly sighted along the trail with an aerial insect net, though many butterflies evaded capture even by experienced surveyors. When captured, butterflies were carefully removed from the net and marked with a unique 3-digit ID number on the ventral surface of both hindwings using a fine-tip permanent marker. Data including sex, time of capture, and location were recorded for each individual captured. For a subset of years (1986 - 2009), forewing length was also measured using digital calipers (Mitutoyo Corp., Illinois, USA). Butterflies were promptly released at the same location where they were captured. Recaptured individuals were identified using the unique ID, and time and location of capture was recorded for each capture event. Butterflies seen but not captured were recorded, but not assigned a unique ID nor used in the analyses presented.

Mark-release-recapture (MRR) models were fit for each year with recaptures in R version 4.2.3 (R Core Team 2020) using the RMark interface (Laake 2013) and Program MARK (White and Burnham 1999). We fit POPAN models, a type of Jolly-Seber model, which themselves are an open population capture-recapture model capable of estimating population size (Schwarz and Arnason 1996). The POPAN Jolly-Seber model estimates four parameters, (1) detection probability (*p_t_*; the probability of detecting an individual at time *t* if it is alive), (2) apparent survival *(√_t_*; the probability of an individual surviving to the next time step), (3) super population size (*N_super_*); the total number of individuals available to enter the study, (4) probability of entry through births and immigration (*pent_t_*; probability of new individuals from the super-population entering at time *t*). We included several potential covariates into estimating the probability of detecting an individual (*p_t_*) to account for detection differences across male and female butterflies, differing expertise of surveyors, and survey effort. These covariates include the sex of the individual butterfly, the primary observer of a survey date, and the total number of butterfly species observed by the surveyors on a survey date (i.e., list length (Breed et al., 2013)). We included the sex of the individual butterfly as a covariate that could impact the estimation of the survival (*t*). We did not include any covariates in the probability of entry submodel (*pent_t_*). Year-specific MRR models were selected based on AICc values (Burnham and Anderson 2004). Two year-specific population estimates had confidence intervals far too large to be informative due to a lack of recapture data and these estimates were removed from further analyses. In total, we had 23 years with population estimates spanning 1986 to 2021.

### 2.4 Weather data

We obtained daily values of minimum and maximum temperature and precipitation values for Elliott Key from 01 January 1985 - 31 December 2021 from Daymet (Thornton et al. 2016). Daily wind speed values came from the second Modern-Era Retrospective analysis for Research and Applications (MERRA-2), a NASA atmospheric reanalysis with data available from 1980 (Global Modeling and Assimilation Office 2015). MERRA-2 estimates surface wind speed (m s^-1^) at two-hour intervals, which we aggregated to the maximum daily wind speed. We included temperature, precipitation, and wind covariates in our modeling framework used to estimate population dynamics.

Given the uncertainty of how weather affects Schaus’ swallowtail (or other tropical butterfly) population dynamics, we used daily weather data to generate a variety of potential predictor variables that were included in exploratory models estimating potential drivers of population dynamics. Year-specific mean minimum temperature and mean maximum temperature values were calculated across three different time periods within a calendar year reflecting the time of year Schaus’ swallowtail were expected to be in different life stages. For each year, we averaged the daily minimum and maximum temperature over key time periods for adult butterflies (day of year [DOY] 121 to 212; i.e., May through July), larvae (DOY 91 - 181; i.e., April through June), and pupae (DOY 213 - 273; i.e., August through September). Year-specific precipitation values summed the daily precipitation accumulated over two time periods representing the wet and dry seasons. The wet season corresponds to May 15 through October 15 (DOY 135 - 288) on a non-leap year. The dry season corresponds to October 16 through May 14 (DOY 289y-1 and 134). For example, the dry season precipitation calculation for the year 2019 would sum the total precipitation accumulated from October 16, 2018 to May 14, 2019.

The two wind variables, maximum wind speed and extreme wind events, were quantified over the entire year. Maximum wind was calculated as the maximum daily wind speed observed in a year, and extreme wind events were calculated as the number of days in which the maximum daily wind speed exceeded 15 m s^-1^, corresponding to frequent gusts that classify as a ‘moderate’ high wind threat level on the National Oceanic and Atmospheric Administration’s wind threat categories (National Weather Service 2023). Temporal lags in species’ response to disturbance and conservation actions are common (Watts et al., 2020), and thus we also included time lags of weather variables as potential predictor variables in downstream modeling. For the temperature and precipitation variables, we included variables with a one-year lag (*t-1*), and for the wind variables, we included variables with up to a 4-year lag (*t-4*) given plant communities may take multiple years to recover following large hurricanes (Rastetter et al. 2021). All predictor variables were scaled to a mean of zero and standard deviation of one to facilitate the interpretation of their relative influence.

### 2.5 Statistical Analysis

#### 2.5.1 Modeling population dynamics

We fit exploratory (sensu Tredennick et al., 2021) linear models to estimate potential drivers of Schaus’ swallowtail population dynamics. In these models, we use the number of survey days as a weight in fitting our regression to give higher weights to years with more sampling effort. Our response variable in these models was the log transformed population estimate generated from the MRR model. Within each class of weather variable (temperature, precipitation, and wind), we fit individual models for each variable and used the Akaike Information Criteria with a correction for small sample size (AICc) to select the best predictors. We retained variables for which AICc was less than 2 and that were not highly correlated (r < 0.6). In the cases where the top univariate predictors were correlated, we chose to retain the variable that was determined to be more likely to influence the population dynamics of the Schaus’ butterfly given expert understanding of the butterfly’s biology (e.g., choosing maximum temperature during larval life stage over maximum temperature during the adult life stage). Following this procedure, we retained the following nine variables: mean maximum temperature of larval stage, mean minimum temperature of the adult stage the year prior (*t*-1), mean maximum temperature of the adult stage the year prior (*t*-1), mean maximum temperature of the pupal stage the year prior (*t*-1), precipitation over the dry season, maximum wind speed, maximum wind speed the year prior (*t*-1), maximum wind speed four years prior (*t*-4) and number of extreme weather events four year priors (*t*-4).

Next, we fit 36 competing models (Table S1), each representing a unique combination of two-variable models from our nine retained predictor variables, to quantify the effect of covariates on the estimated population size. We included a maximum of two predictor variables in our competing models to adhere to the 10 data points per predictor rule, since we were limited to 23 years with a population estimate. We calculated the relative importance of each variable by summing the AICc weights for each model in which that variable appears (Symonds and Moussalli, 2011). For a subset of competing models with AICc ≤ 2, we used AICc weighted averaging (Burnham and Anderson 2004) to calculate model averaged coefficient estimates and confidence intervals. Residual diagnostic tests of the model with the lowest AICc were conducted to ensure that model assumptions were met (Figure S1). Residual diagnostic tests of the model with the lowest AICc were conducted to ensure that model assumptions were met (Figure S1).

We also fit models to test the importance of density dependence on Schaus’ swallowtail population dynamics. To do so, we first identified years with consecutive population estimates, which allowed us to include a population estimate of the prior year as a predictor variable. Given this data structure, we calculated a population growth rate parameter, instantaneous rate of population increase 𝑟 = (𝑙𝑜𝑔𝑁_1_/𝑁_0_); when *N_0_* = population size at time zero and *N_1_* = population size at time 1), which we used as the response variable for this set of models. In total we had 14 years with a population estimate that was also estimated for the year prior. Here, we built 15 competing two-variable models predicting population growth rate based on the five variables retained in the top population estimate models described above and also including the new density dependence predictor variable (log[population estimate at *t*-1]; Table S2). As detailed above, we calculated the relative importance of each variable across all models, and the model-averaged coefficient estimates and confidence intervals for competing models with AICc ≤ 2. Residual diagnostic tests of the model with the lowest AICc were conducted to ensure that model assumptions were met (Figure S2).

#### 2.5.2 Modeling forewing size

To test if forewing size was influenced by weather conditions, we built a linear model with forewing size as a response variable using the following two continuous predictors: mean maximum temperature during the larval time period and precipitation during the wet season in the year prior. We also included butterfly sex as a categorical predictor.

## 3. Results

### 3.1 Population dynamics

The population estimates of the Schaus’ swallowtail varied considerably over time, with the smallest population size estimated for the year 2007 (56 [95% CI 41 - 77]), and the largest population size estimated for the year 1987 (11,360 [95% CI 4,505 - 28,673]. The population estimates did not show a consistent positive or negative trend through time (Figure 2A). The probability of capturing an individual alive at any time (*t*) was higher for males (median *p_t_* value of 0.126 across all years) than for females (median *p_t_* value of 0.072), and the probability an individual survived one day was also higher for males (median √ value of 0.738) than females (median √ value of 0.653).

**Figure 2.**
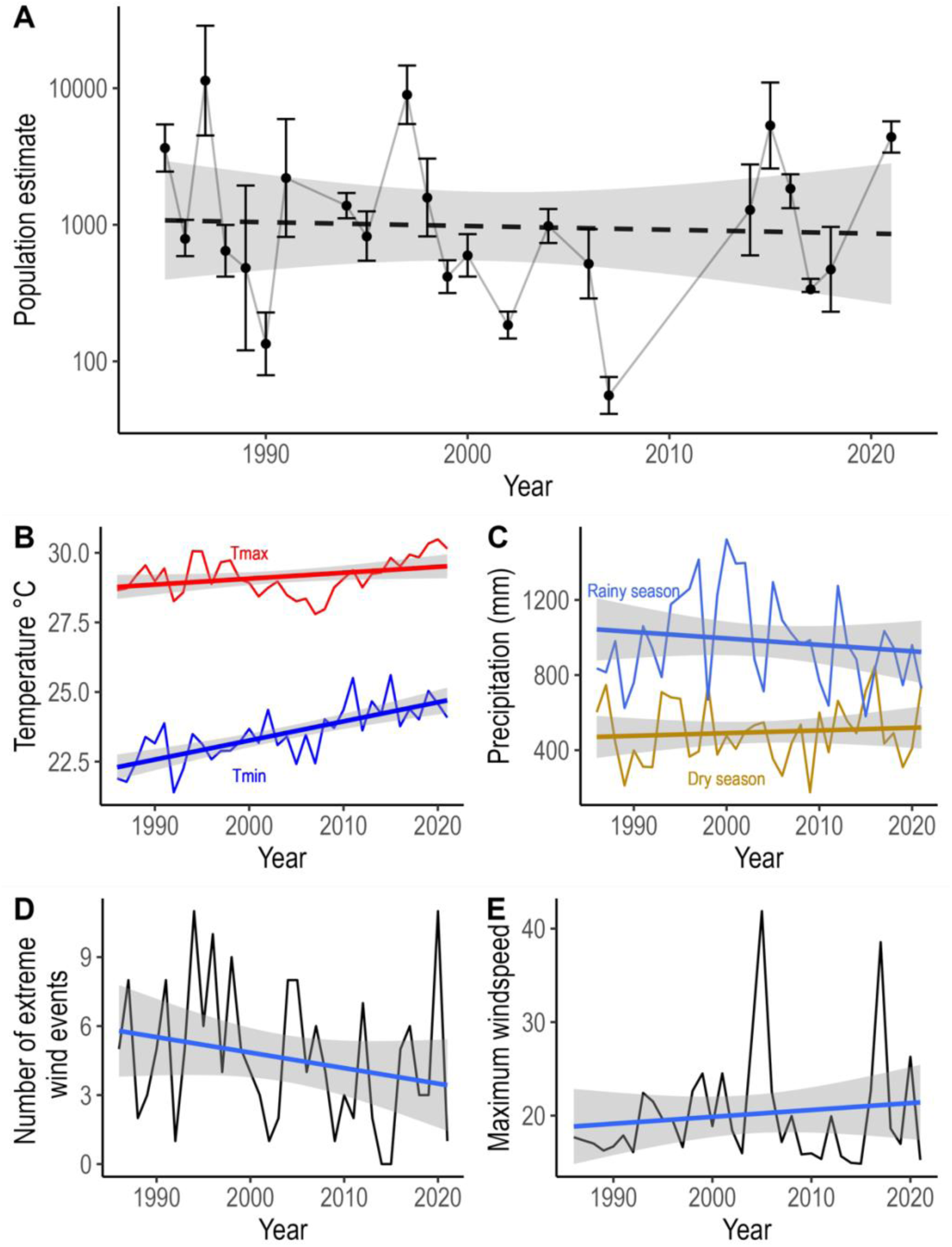
Variation of Schaus’ swallowtail population estimates and weather variables from 1985-2021. Schaus’ swallowtail population estimates show large variation across years, but no significant trend (A). Maximum temperature during the larval stage has remained relatively consistent across our study period (red), but minimum temperature has been increasing (blue) (B). Wet season (blue) and dry season (bronze) annual precipitation (C), number of extreme wind events (D), and maximum wind speed (E) had large interannual variation but no long-term trend.

The most important weather variables predicting the variation in population size were wind variables. We found an 80% chance that a wind variable is a component of the best model among our competing models (Figure 3A). Wind variables affected population size differentially depending on the temporal lag since the wind event occurred. Both maximum wind in the year and maximum wind in the previous year negatively impacted Schaus swallowtail population size, while population size was higher four years after a year with high winds (Figure 3B). Precipitation variables were the second most important weather variables predicting population size, with population size being larger in years with more rain during the dry season (i.e. the season prior to adult emergence) (Figure 3B). Temperature variables were the least important weather variables predicting population size with the probability that a temperature variable is a component of the best model among our competing models being 40% (Figure 3A). Warmer minimum temperatures during the adult flight season the year prior led (*t-1*) to smaller population sizes in a given year (*t*) (Figure 3B).

**Figure 3.**
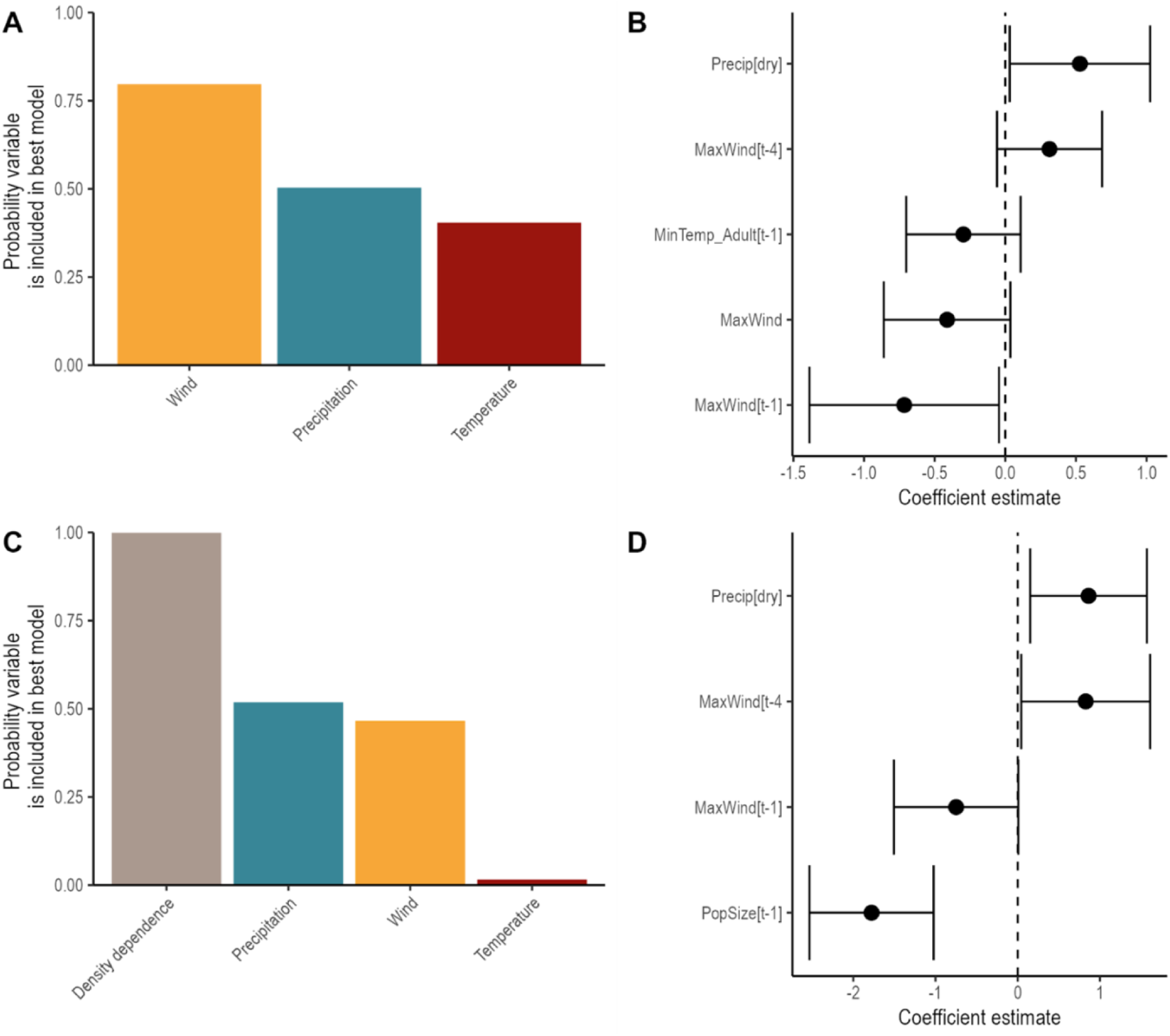
Results of exploratory models estimating po pulation estimate (A,B) and growth rate (C,D). Wind variables are most likely to be included in a best model estimating population size (A), while density dependence is most likely to be included in a best model estimating growth rate (C). Model averaged coefficients show population size increases in years with more rain during the dry season and decreases two years after years with heavy winds (B). Growth rates of Schaus’ swallowtails display negative density dependence (D).

The most important variable predicting variation in growth rate of Schaus’ swallowtail population size on Elliott Key was the population size the year prior (i.e., density-dependence) with a 99.9% probability of density-dependence being a component of the best model among our competing models (Figure 3C). The next two most important variables were precipitation (51.8% probability) followed by wind (46.6% probability). It was unlikely temperature variables were important to explaining the variation in population growth rate (1.6% probability). The model averaged coefficient estimates of the variables retained in the models with an AICc ≤ 2 shows that population growth rate from one year to the next was lower in years when population size was higher the previous year, indicating negative density-dependence (Figure 3D). Growth rate was also lower in years following a year with high winds. In contrast, it was higher in years when precipitation was high during the dry season and four years after a year with high winds (Figure 3D).

### 3.2 Variation in forewing size

Sex, temperature, and precipitation were all important in estimating variation in Schaus’ swallowtail forewing length (Table S3). Schaus’ swallowtails are sexually dimorphic; female butterflies are larger than males (male coefficient estimate = -4.17 [95% CI -4.38 – -3.95]; Figure 4A). Our model estimating forewing length of individual butterflies found that forewings were smaller in years for which the larvae experienced warm temperatures (coefficient estimate -0.29 [95% CI -0.38 – -0.20]; Figure 4B). Our model results also estimate forewing length to be larger in years following with more precipitation in the wet season the year prior (coefficient estimate 0.63 [95% CI 0.54 – 0.73]; Figure 4C). Overall, sex was the most important variable in explaining variation in forewing length, while weather variables explained far less variation in forewing length (R^2^ = 0.03 in a two variable model that does not include sex). Together, the three variable model had an R^2^ = 0.37.

**Figure 4.**
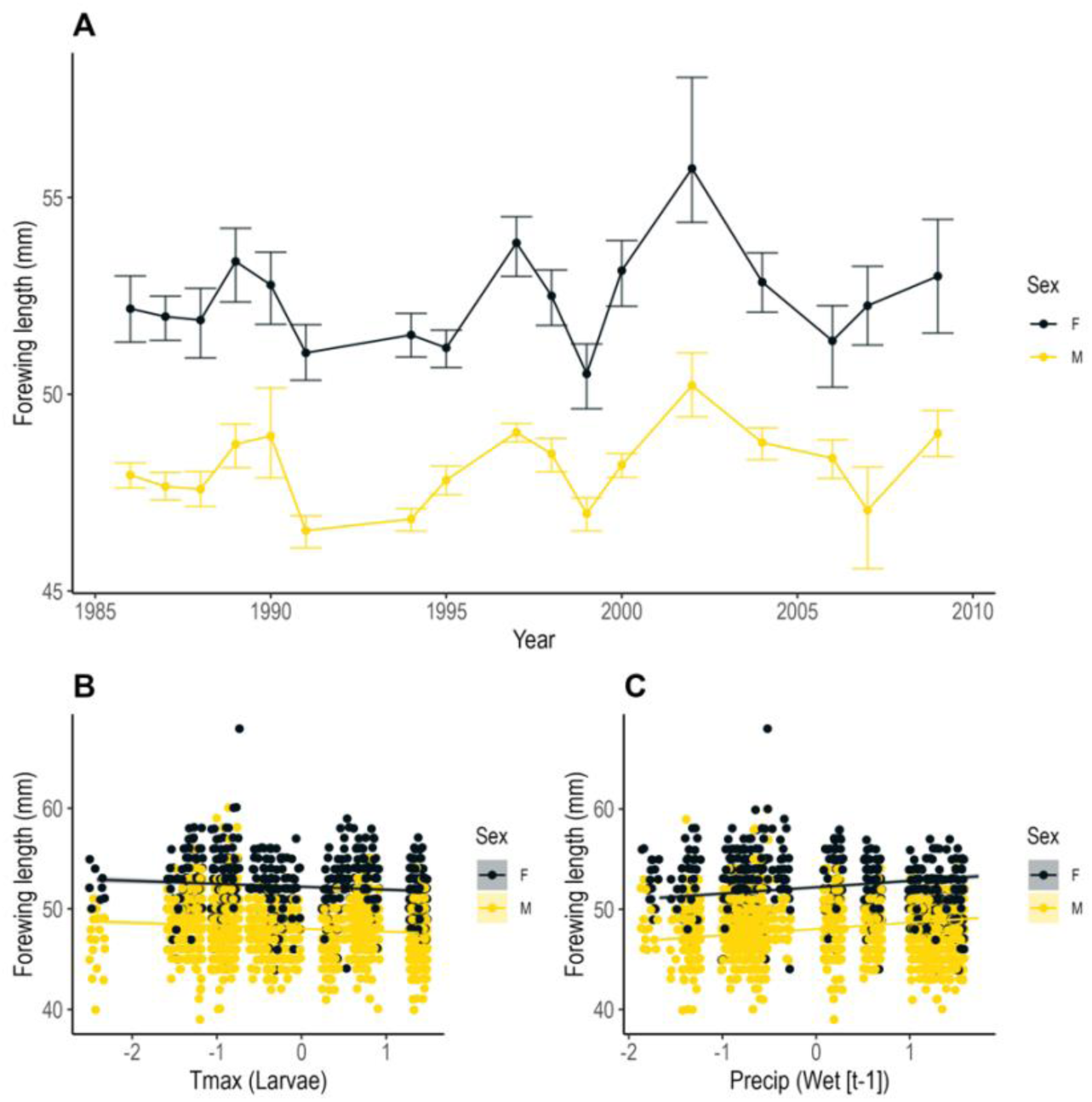
Measurement of Schaus’ swallowtail forewing length (and 95% bootstrapped CIs) for a subset of study years (A). Model predictions show forewing length was smaller following warmer years during the adult larval stage (B), but forewing length was larger in years with more rain in the wet season the year prior (C). Females were on average larger than males. Points are jittered to show density of data.

## 4. Discussion

The endangered Schaus’ swallowtail butterfly is a flagship of insect conservation, as it was among the first butterfly species to be listed as threatened under the Endangered Species Act in the United States and is a key umbrella species for tropical hardwood hammocks in Florida. Unfortunately, this butterfly is still at high risk of extinction due to habitat destruction, climate change, and sea level rise. Our study leverages a remarkable dataset, likely the longest running mark-recapture dataset of any tropical butterfly, to provide ecological insights into species biology and population dynamics. We provide quantitative evidence of long-held speculations, including that females are larger than males, and uncover a complex response of Schaus’ swallowtail butterfly to hurricane-mediated disturbance. Specifically, we report evidence that high winds associated with tropical cyclones immediately decrease Schaus’ swallowtail population size, but the longer-term disturbance leads to increases in population size several years following a tropical cyclone event.

The 35-year dataset reveals that this population of Schaus’ swallowtail butterfly is stable over the long term, which was only revealed due to the long-term nature of this study. Conclusions regarding population status and trend may have differed significantly had the study only been conducted for a subset of the whole dataset. In the short term, population size was highly variable, with population estimates ranging from under 100 individuals to over 10,000 individuals. Basing conclusions on only a few years would demonstrate ‘the snapshot effect’ in which population trend estimates can be greatly influenced by the choice of contemporary time-points (Didham et al. 2020). We echo the calls of others (Dolný et al. 2021; Hellmann et al. 2003) to interpret the results of population trends using short time-series data, especially for insects, with caution given the limited statistical power of such data.

This study demonstrates that the Elliott Key population of Schaus’ swallowtail butterfly is highly resilient to both environmental and demographic stochasticity. Population dynamics are most influenced by population size in the previous year, followed by high wind events and precipitation. Schaus’ swallowtail population dynamics are associated with recent high wind events and precipitation, two prominent features of tropical cyclones. This is perhaps unsurprising since this butterfly is endemic to South Florida, a region that experiences frequent hurricanes, with a major hurricane approximately every 30 years (Doyle and Girod 1997). During the study period analyzed, dozens of hurricanes tracked through southeastern Florida, with the most notable being Hurricane Andrew, a compact Category 5 storm that made landfall in August 1992 (National Oceanic and Atmospheric Administration 2024). Hurricane Andrew was a devastating storm for Biscayne National Park, with virtually all large trees located in hardwood hammock forests defoliated and about 25% windthrown or badly broken (NPS 2019). Despite the enormous magnitude of this disturbance and likely mortality of butterflies due directly to high winds and storm surge as well as decreased resources in the months following Hurricane Andrew, population numbers rebounded only a few years later.

One mechanism which could explain an increase in the population of Schaus’ swallowtail butterfly several years following a hurricane is an increase in floral resources that result in increased nectar availability for adult butterflies. It is well-established that the high wind and heavy precipitation associated with tropical cyclones create treefall gaps which can result in higher local plant diversity (Lugo 2008; Mitchell 2013; Murphy et al. 2014; Shiels et al. 2015). In the 2-3 years after Hurricane Andrew at nearby study sites in Everglades National Park and elsewhere in the Upper Florida Keys, early successional woody and herbaceous species dominated previously closed-canopy hardwood hammock forests and understory trees flowered much more extensively than prior to the hurricane (Armentano 1995; Ross et al. 2001). Although the impact of hurricanes on plant flowering and fruiting phenology is highly variable (Rathcke 2000; Gandhi et al. 2007), one study showed an increase in flowering of a common understory shrub in southeastern Florida 2-3 years after Hurricane Andrew in hurricane-damaged populations, while nearby undamaged populations did not flower at all (Pascarella 1998). Increased nectar resources result in increasing butterfly survivorship and fecundity (Hill and Pierce 1989; Boggs and Ross 1993; Romeis et al. 2005); nectar can be a limited resource in nature (Schultz and Dlugosch 1999; Erhardt and Mevi-Schütz 2009). We believe this could be the case in the closed canopy forest habitat of Schaus’ swallowtail butterfly, where butterflies are frequently observed nectaring on the flowers of herbaceous plants along trails, treefall gaps, and other clearings with high light availability. Population decreases the year following high winds is likely due to increased mortality across all life stages from storm surge inundation and vegetation defoliation from high winds which decrease oviposition habitat and larval resources. Negative density dependence is common but not ubiquitous among Lepidoptera and other insects (Dempster 1983; Stiling 1988). In this case, negative density dependence could be explained by intraspecific competition (Nowicki et al. 2009; Flockhart et al 2012) or parasitism (Daniels et al. 1993).

As we demonstrate here with Schaus’ swallowtail butterfly, taxa in the Caribbean Basin are well-adapted to persist with frequent hurricane events (Schowalter et al. 2021). Global climate change is projected to increase the intensity of hurricane events in the Atlantic Basic (Bender et al. 2010) and sea level rise will exacerbate storm surge risk (Walsh et al. 2016). Although Schaus’ swallowtail butterfly and other organisms that inhabit this region are clearly well-adapted to be resilient to tropical cyclone events (Donihue et al. 2018), ecological disturbance events can increase the risk of extinction for taxa with small and fragmented populations (Casagrandi and Gatto 2002; Ovaskainen and Meerson 2010), particularly narrow-range endemics and taxa are already endangered due to human factors (Goulding et al. 2016; Crain et al. 2019). This seems to be the case for endemic Caribbean birds such as the Puerto Rican parrot (*Amazona vittata*), which became endangered due to habitat destruction and harvesting of birds for pets and food (Snyder et al. 1987). Although captive breeding and reintroduction efforts have successfully increased this number, severe hurricanes periodically cause massive mortality of wild parrots (Beissinger et al. 2008; Martínez and Logue 2020).

This study provides valuable insights into the natural and life history of Schaus’ swallowtail and substantiates many anecdotal observations made by researchers. We confirm sexual dimorphism in this butterfly, with females having a consistently larger forewing length than males. This female-biased sexual size dimorphism is common in arthropods and is generally attributed to the greater fecundity of larger females (Shine 1989; Head 1995; Teder and Tammaru 2005). We also confirm the presence of adult butterflies in August and September. Although data were insufficient to understand population size of this second flight period, these observations raise the question of whether this butterfly could have two generations per year rather than one generation per year, as previously thought. Further exploration of the prevalence of a secondary flight period is warranted for this taxon.

Across the study period, males were nearly twice as likely to be captured as females, and the probability of recapturing males was also higher than the probability of recapturing females. This finding, combined with anecdotal observations of apparent patrolling behavior along the trail by males with individual males often sighted repeatedly at the same location, indicates that male Schaus’ swallowtail butterflies are likely territorial; male territoriality has been observed in several other papilionid species (Lederhouse 1982; Pinheiro 1990; Lehnert et al. 2013). We conjecture that males may be patrolling the increased nectar flower resources available along canopy openings such as trails, which are attractive to foraging females, making males them more likely to be encountered by researchers than females. When encountered, females often appeared to be searching for or evaluating host plants for oviposition. Since the primary host plant for Schaus’ swallowtail butterfly is an understory tree, it is found throughout the habitat in areas that are far from the trail and difficult to access due to the thick vegetative growth which may account for the sex bias in captures.

As is common with data collected over multiple decades, survey effort was variable and dependent upon fluctuating funding. This resulted in some years with large confidence intervals surrounding population estimates and years for which population size could not be estimated. Across the entire study period, there were dozens of observers with varying levels of experience which led to differing capture rates between observers. Additionally, this butterfly is the subject of past and ongoing efforts to restore populations through releases of individuals head started under laboratory conditions.

During 1995-97, over 1000 butterflies were released within Biscayne National Park, North Key Largo, Miami, and other locations within the historical range for Schaus’ swallowtail. In 2014-21, over 1300 individuals were released in Biscayne National Park and nearby areas. To support these releases, several dozen individuals, usually larvae, were removed from the wild in the year prior to releases. We use an open population model to account for migration, but these releases may represent an additional source of uncertainty in population estimates. Many of the head started butterflies were released during the larval stage which are subject to a high rate of predation (Clayborn and Koptur 2017), so the number of head started individuals that contributed to population estimates is likely much smaller than the total number released.

The persistence of the Schaus’ swallowtail population on Elliott Key highlights the success of past and ongoing efforts to conserve and manage remaining butterfly populations and hardwood hammock forests within their range – critical conservation efforts particularly given the destruction of much of this habitat in the 20th century (Walker et al. 1997). Our study site, Elliott Key, supports what is likely the most robust population of Schaus’ swallowtail across its range. Although population estimates on other islands within Biscayne National Park and on Key Largo have not been conducted, community science surveys of Key Largo indicate that Schaus’ swallowtail’s main host plant is less abundant than on Elliott Key and numbers of butterflies observed during Pollard transects are indicative of smaller populations in Key Largo than Elliott Key (unpublished data). This unprecedented continuous dataset spanning five decades not only sheds light on the natural history and ecology of a charismatic and endangered insect; it also permits us to examine truly long-term population trends. The nearly four-decade dataset allows us to see the volatility of population dynamics while contextualizing them as that - volatility around a long-term stability and not true trends. This study underscores the role that tropical cyclone disturbance plays in species population dynamics, highlighting the need for continued efforts to document its role in maintaining tropical diversity.

## Acknowledgements

This publication represents nearly 40 years of diligent fieldwork conducted in extremely challenging field conditions. It takes true dedication to don a mosquito jacket in the heat of summer and eschew bug spray to chase rare butterflies! Over the years, dozens of people have contributed to these field survey efforts, including but not limited to: Lukasz Barszczak, Jonathan Bremer, Edson Bustamente, Keith Curry-Pochy, Peter J. Eliazar, Ryan Fessenden, Mark Goode, Leslie G. Groce, B. J. Haeck, Geena M. Hill, Taylor S. Hunt, Chase B. Kimmel, Shawn D. Larson, Marc C. Minno, Ray Moranz, James Nation, Jr., J. Akers Pence, Lary Reeves, Arif Rehman, David B. Ritland, Erik Runquist, Mark Salvato, Stephanie Sanchez, Jamie Sarvis, James B. Schlachta, Steven Schlachta, Andrei Sourakov, Kevina Vulinec, Samm Wehmen Epstein, and Keith Willmott Staff at the McGuire Center for Lepidoptera & Biodiversity, particularly Charles V. Covell Jr. (then of University of Louisville, Kentucky), Christine Eliazer, Peter J. Eliazar, were critical to ensuring survey continuity and long-term support. Collaborators and staff at Biscayne National Park, especially Elsa Alvear, Richard Curry, and Vanessa McDonough, were fundamental to securing access to sites, which is no small task for an uninhabited island and involved a great deal of boat rides and trail clearing efforts.

Finally, staff at U.S. Fish and Wildlife Service, particularly Mark Salvato, were indispensable in securing permits and providing other support throughout the years. We also wish to thank Jesse Borden, Phil Hahn, and Laura Steele for their helpful feedback on a draft of this manuscript.

