## Supplementary material for "Long-term population dynamics of an endangered butterfly are influenced by hurricane-mediated disturbance"

Table S1. Summary of all models fitted to predict population dynamics of Schaus’ swallowtail. Included predictor variables are shown in the Model column.

| Model | ∆AICc | Weight |
| --- | --- | --- |
| MaxWind + Precip[dry] | 0.000 | 0.144 |
| MaxWind[t-4] + Precip[dry] | 0.749 | 0.099 |
| MaxWind + MaxWind[t-1] | 0.792 | 0.097 |
| Precip[dry] + MinTempAdult[t-1] | 1.233 | 0.078 |
| MaxWind + MaxWind[t-4] | 1.929 | 0.055 |
| Precip[dry] + MaxTempAdult[t-1] | 2.030 | 0.052 |
| ExtremeWind[t-4] + MaxWind | 2.255 | 0.047 |
| ExtremeWind[t-4] + Precip[dry] | 2.318 | 0.045 |
| Precip[dry] + MaxTempLarvae | 2.837 | 0.035 |
| MaxWind[t-1] + MaxWind[t-4] | 2.839 | 0.035 |
| ExtremeWind[t-4] + MaxTempLarvae | 2.882 | 0.034 |
| ExtremeWind[t-4] + MaxWind[t-4] | 3.160 | 0.030 |
| MaxWind[t-1] + Precip[dry] | 3.300 | 0.028 |
| MaxWind + MaxTempLarvae | 3.523 | 0.025 |
| MaxWind[t-4] + MaxTempLarvae | 3.567 | 0.024 |
| Precip[dry] + MaxTempPupae[t-1] | 3.765 | 0.022 |
| MaxWind[t-4] + MaxTempAdult[t-1] | 3.875 | 0.021 |
| ExtremeWind[t-4] + MaxWind[t-1] | 4.433 | 0.016 |
| MaxWind[t-4] + MinTempAdult[t-1] | 4.443 | 0.016 |
| MaxWind[t-4] + MaxTempPupae[t-1] | 4.517 | 0.015 |
| ExtremeWind[t-4] + MaxTempPupae[t-1] | 4.829 | 0.013 |
| MaxWind + MinTempAdult[t-1] | 5.822 | 0.008 |
| MaxWind + MaxTempAdult[t-1] | 5.831 | 0.008 |
| MaxWind + MaxTempPupae[t-1] | 5.858 | 0.008 |
| ExtremeWind[t-4] + MinTempAdult[t-1] | 6.218 | 0.006 |
| ExtremeWind[t-4] + MaxTempAdult[t-1] | 6.232 | 0.006 |
| MaxWind[t-1] + MaxTempLarvae | 6.802 | 0.005 |
| MaxWind[t-1] + MaxTempAdult[t-1] | 6.809 | 0.005 |
| MaxTempAdult[t-1] + MaxTempLarvae | 6.818 | 0.005 |
| MaxWind[t-1] + MinTempAdult[t-1] | 6.834 | 0.005 |
| MaxTempLarvae + MinTempAdult[t-1] | 7.075 | 0.004 |
| MaxWind[t-1] + MaxTempPupae[t-1] | 7.509 | 0.003 |
| MaxTempLarvae + MaxTempPupae[t-1] | 8.114 | 0.002 |
| MaxTempPupae[t-1] + MinTempAdult[t-1] | 8.830 | 0.002 |
| MaxTempAdult[t-1] + MaxTempPupae[t-1] | 8.940 | 0.002 |
| MaxTempAdult[t-1] + MinTempAdult[t-1] | 9.422 | 0.001 |

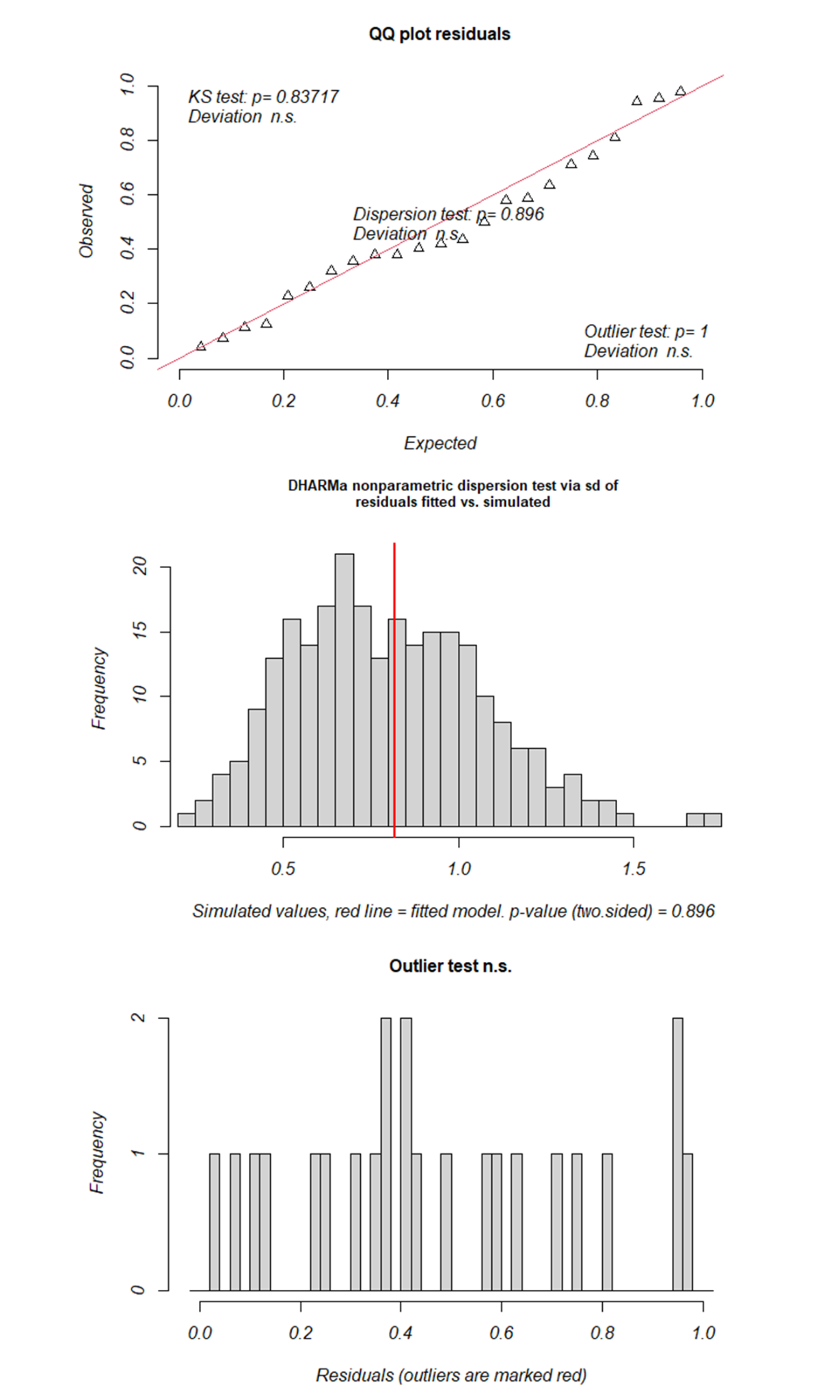

Figure S1. Summary of residual diagnostic test for population dynamics model with the lowest ∆AICc.

Table S2. Summary of all models fitted to predict growth rate of Schaus’ swallowtail. Included predictor variables are shown in the Model column.

| Model | delta | weight |
| --- | --- | --- |
| PopSize[t-1] + Precip[dry] | 0.000 | 0.518 |
| PopSize[t-1] + MaxWind[t-4] | 1.418 | 0.255 |
| PopSize[t-1] + MaxWind[t-1] | 1.966 | 0.194 |
| PopSize[t-1] + MaxWind | 6.858 | 0.017 |
| PopSize[t-1] + MinTempAdult[t-1] | 6.997 | 0.016 |
| MaxWind + MaxWind[t-1] | 15.248 | 0.000 |
| MaxWind + Precip[dry] | 16.383 | 0.000 |
| Precip[dry] + MinTempAdult[t-1] | 17.056 | 0.000 |
| MaxWind[t-1] + Precip[dry] | 17.099 | 0.000 |
| MaxWind + MaxWind[t-4] | 17.773 | 0.000 |
| MaxWind[t-1] + MinTempAdult[t-1] | 17.781 | 0.000 |
| MaxWind + MinTempAdult[t-1] | 17.783 | 0.000 |
| MaxWind[t-4] + Precip[dry] | 18.223 | 0.000 |
| MaxWind[t-1] + MaxWind[t-4] | 18.240 | 0.000 |
| MaxWind[t-4] + MinTempAdult[t-1] | 19.148 | 0.000 |

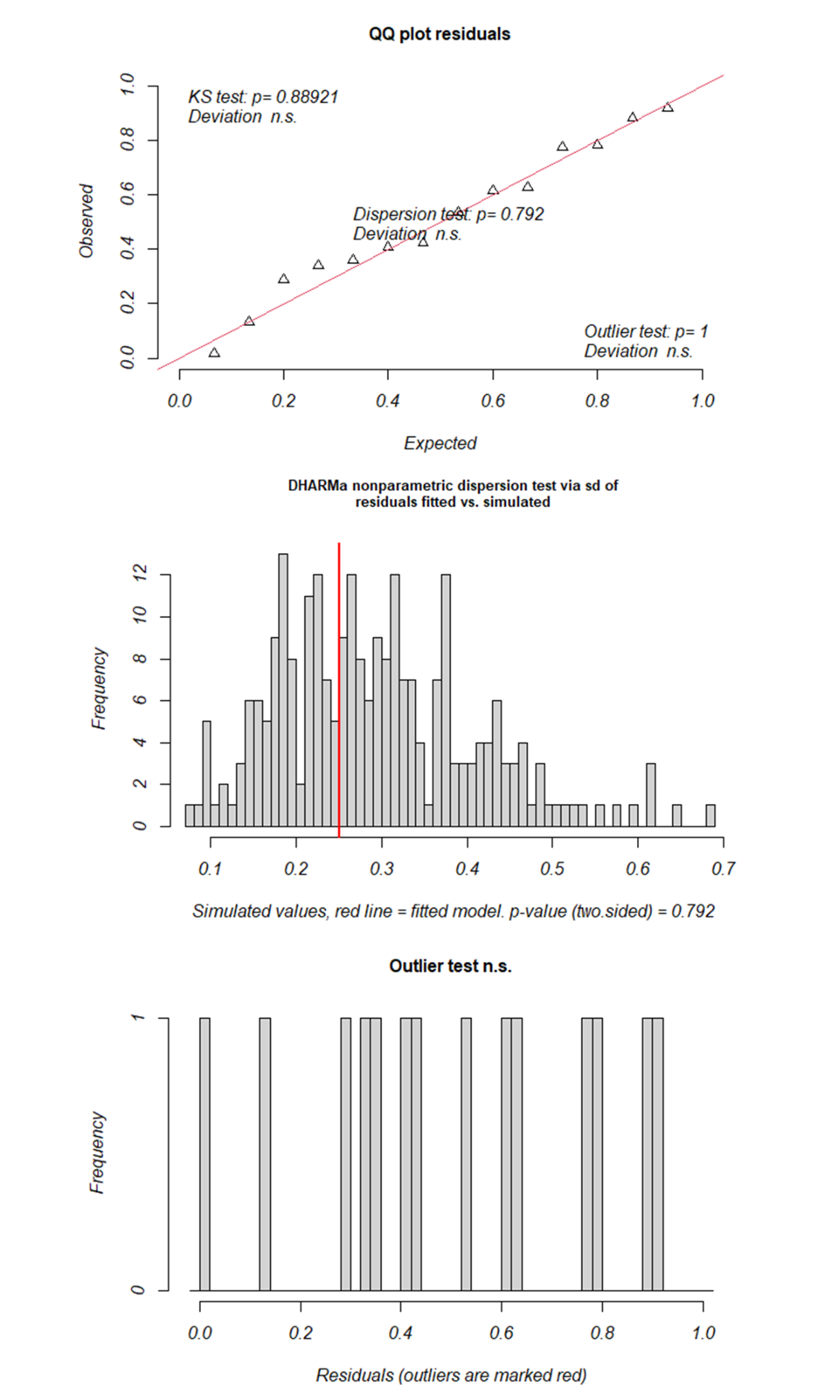

Figure S2. Summary of residual diagnostic tests for the population growth rate model with the

lowest ∆AICc.

Table S3. Coefficient estimates for model estimating forewing length.

| Model Coefficients | Estimate | 2.5% CI | 97.5% CI |
| --- | --- | --- | --- |
| (Intercept) | 52.19 | 52.01 | 52.38 |
| Tmax[Larvae] | -0.29 | -0.39 | -0.20 |
| Prcp[Rainy] (t-1) | 0.63 | 0.54 | 0.73 |
| Sex[M] | -4.17 | -4.39 | -3.96 |
